# Antibody response to SARS-CoV-2 mRNA vaccines in pregnant women and their neonates

**DOI:** 10.1101/2021.04.05.438524

**Authors:** Malavika Prabhu, Elisabeth A. Murphy, Ashley C. Sukhu, Jim Yee, Sunidhi Singh, Dorothy Eng, Zhen Zhao, Laura E. Riley, Yawei J. Yang

**Affiliations:** Department of Obstetrics and Gynecology, Weill Cornell Medicine, New York, NY, USA; Laboratory of Molecular Neuro-oncology, The Rockefeller University, New York, NY, USA; Department of Pathology, Weill Cornell Medicine, New York, NY, USA; Department of Pathology, New York-Presbyterian Hospital/Weill Cornell Medical Center, New York, USA

## Abstract

Pregnant women were excluded from initial clinical trials for COVID-19 vaccines^1–2^, thus the immunologic response to vaccination in pregnancy and the transplacental transfer of maternal antibodies are just beginning to be studied^4–5^.

## Methods

Between January 28 and March 31, 2021, we studied 122 pregnant women and their neonates at time of birth. All women had received one or both doses of a messenger RNA (mRNA)-based COVID-19 vaccine. Fifty-five women received only one dose of the vaccine and 67 women received both doses of the vaccine by time of giving birth. Eighty-five women received the Pfizer-BioNTech vaccine, while 37 women received the Moderna vaccine. All women tested negative for SARS-CoV-2 infection using reverse-transcriptase PCR on nasopharyngeal swabs, and none reported any COVID-19 symptoms at the time of admission for birth. Semi-quantitative testing for antibodies against S-Receptor Binding Domain (RBD) (ET HealthCare)^3^ was performed on sera of maternal peripheral blood and neonatal cord blood at the time of delivery to identify antibodies mounted against the vaccine. All women tested negative for antibodies against the Nucleocapsid Protein (NP) antigen (Roche Diagnostics EUA) to ensure that the antibodies detected were not produced in response to past SARS-CoV-2 infection.

Relationship between IgG antibody levels over time was studied using ANOVA with Tukey posthoc. Relationship between maternal and neonatal IgG levels was studied using Pearson correlation analysis and linear regression on log2-scaled serological values. Relationship between IgG placental transfer ratio (neonatal/maternal) vs. time was studied using Pearson correlation analysis and linear regression on log2-scaled serological values and days. Serology levels represented as log2+1. Statistical analysis was performed using R 3.6.3, RStudio 1.1.463. The study was approved by the Weill Cornell Medicine institutional review board.

## Results

Pregnant women vaccinated with mRNA-based COVID-19 vaccines during pregnancy and tested at time of birth had detectable immunoglobulin (Ig)G and IgM response. Eighty-seven women tested at birth produced only an IgG response, and 19 women produced both an IgM and IgG response. Sixteen women tested at birth had no detectable antibody response, and they were all within four weeks after vaccination dose 1 (Figure 1A). There was an increase over time in the number of women that mounted an antibody response, as well as the number of women that demonstrated passive immunity to their neonates (Figure 1A). All women and their neonates, except for one neonate, had detectable IgG antibodies by 4 weeks after maternal first dose of vaccination (Figure 1A). 43.6% (24/55) of neonates born to women that received only one vaccine dose had detectable IgG, while 98.5% (65/67) of neonates born to women that received both vaccine doses had detectable IgG.

**Figure 1:**
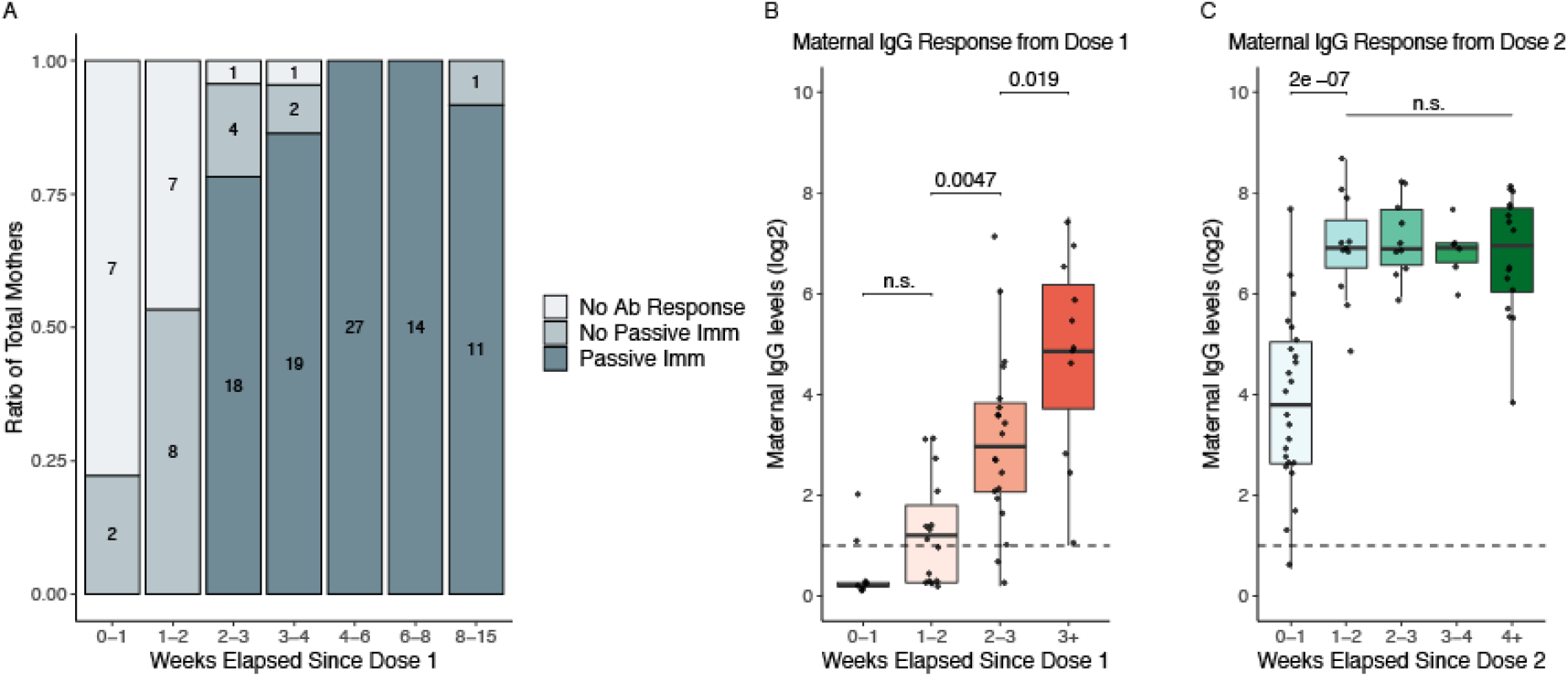
Maternal Antibody Response to Maternal SARS-CoV-2 mRNA Vaccination. **A**. Ratio of women that produced no antibody response (No Ab Response-grey), produced an antibody response but did not demonstrate passive immunity to their neonates (No Passive Imm-light blue), or produced an antibody response and also demonstrated passive immunity to their neonates (Passive Imm-dark blue). Time 0 is day of maternal vaccination dose 1. The earliest detection of antibodies in mothers was at 5 days post maternal dose 1. The earliest detection of antibodies in neonates (passive immunity) was at 16 days post maternal dose 1. The one mother-baby dyad with no transfer of antibodies to the neonate was at 10.14 weeks from dose 1 and 6 weeks from dose 2. **B**. Maternal IgG levels vs. weeks elapsed since maternal vaccination dose 1 for women that only received one dose of the vaccine (55 women). Time point 0 is day of vaccination dose 1. **C.** Maternal IgG levels vs. weeks elapsed since maternal vaccination dose 2 for women that received both doses of the vaccine (67 women). Time point 0 is day of vaccination dose 2. All positive serology cutoffs were 1 (dashed line). Women received either the Moderna or the Pfizer-BioNTech vaccines.

The IgG levels in pregnant women increased weekly from two weeks after first vaccine dose (p=0.0047;0.019), as well as between the first and second weeks after the second vaccine dose (p=2e-07) (Figure 1B). Maternal IgG levels were linearly associated with neonatal IgG levels (R=0.89, p<2.2e-16) (Figure 2A). Placental transfer ratio correlated with the weeks that elapsed since maternal second dose of vaccine (R=0.8, p=2.6e-15) (Figure 2B).

**Figure 2.**
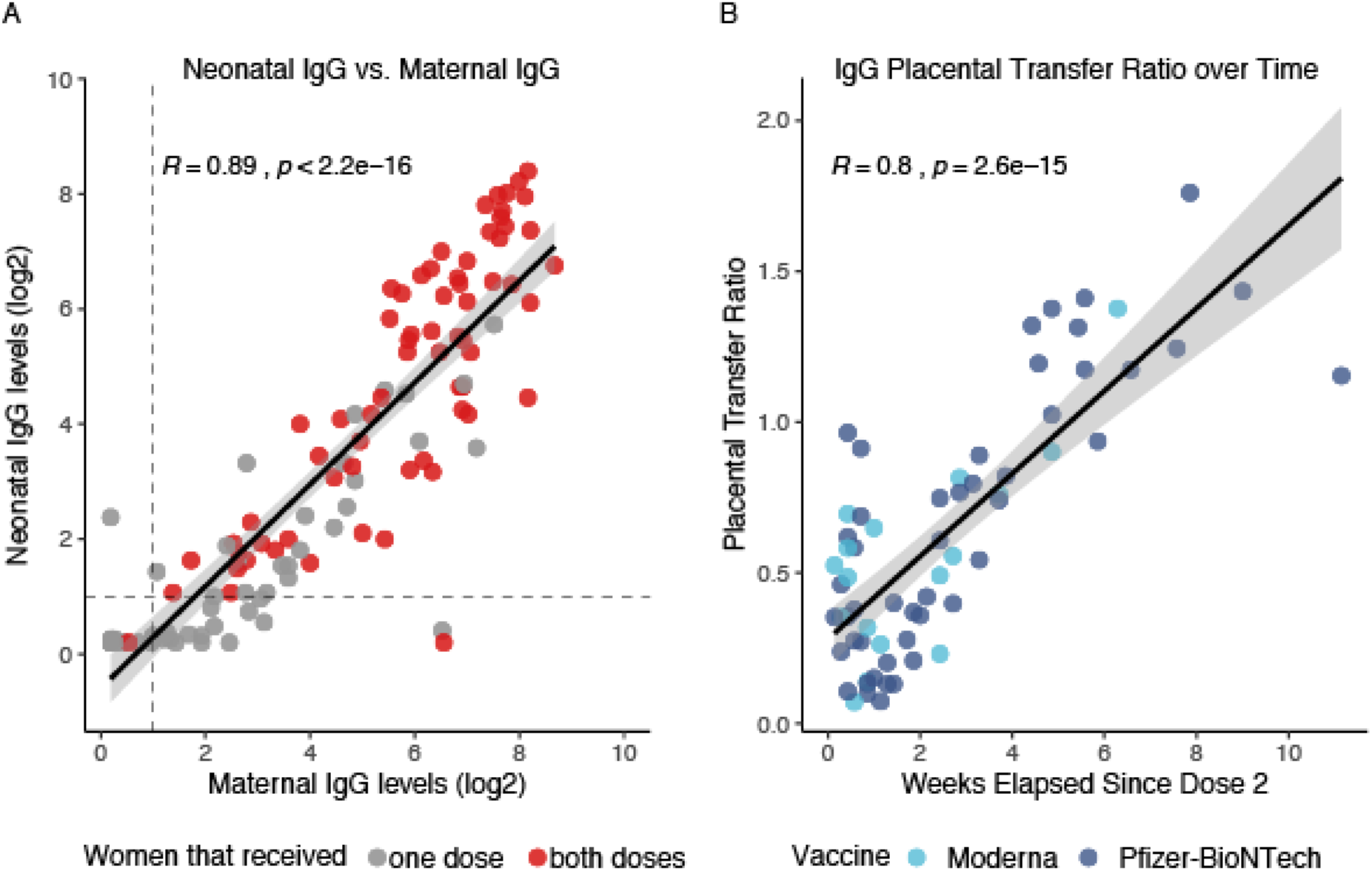
Neonatal Antibody Response to Maternal SARS-CoV-2 mRNA Vaccination. **A.** Neonatal IgG levels vs. maternal IgG levels. Grey dots represent neonates born to mothers that only received one dose of the vaccine. Red dots represent neonates born to mothers that received both doses of the vaccine. All positive serology cutoffs were 1 (dashed line). **B.** Placental transfer ratio (Neonate IgG / Maternal IgG) vs. weeks elapsed since maternal vaccination dose 2 for 65 mother-baby dyads containing mothers that received both vaccine doses. Time point 0 is day of vaccine dose 2. Light blue dots represent maternal Moderna vaccination, dark blue dots represent maternal Pfizer-BioNTech vaccination. Women received either the Moderna or the Pfizer-BioNTech vaccines.

## Discussion

mRNA-based COVID-19 vaccines in pregnant women lead to maternal antibody production as early as 5 days after the first vaccination dose, and passive immunity to the neonate as early as 16 days after the first vaccination dose. The increasing levels of maternal IgG over time, and the increasing placental IgG transfer ratio over time suggest that timing between vaccination and birth may be an important factor to consider in the vaccination strategies of pregnant women. Further studies are needed to understand the factors that influence transplacental transfer of IgG antibody, as well as the protective nature of these antibodies.

## Author Contributions

Dr Yang had full access to all of the data in the study and takes responsibility for the integrity of the data and the accuracy of the data analysis. Concept and design: Prabhu, Riley, Yang.

Acquisition, analysis, or interpretation of data: All authors.

Drafting of the manuscript: Prabhu, Riley, Yang

Critical revision of the manuscript for important intellectual content: All authors. Statistical analysis: Prabhu, Murphy, Yang

Obtained funding: Yang

Administrative, technical, or material support: Prabhu, Sukhu, Yee, Singh, Yang Supervision: Yang

## Additional Contributions

We thank the women of this study, the care staff of Labor & Delivery, and the staff and leadership of Central Laboratories and Blood Bank in the Department of Pathology and Laboratory Medicine for their help in making this study possible. Project support for this research was provided in part by the Center for Translational Pathology at the Department of Pathology and Laboratory Medicine, Weill Cornell Medicine.

